# Global state of deworming coverage and inequity in low-income and middle-income countries: a spatiotemporal study of household health surveys

**DOI:** 10.1101/589127

**Authors:** Nathan C. Lo, Sam Heft-Neal, Jean T. Coulibaly, Leslie Leonard, Eran Bendavid, David G. Addiss

## Abstract

**Introduction:** Mass deworming against soil-transmitted helminthiasis (STH) is a hallmark program in the neglected tropical diseases portfolio that is designed to be equitable and “pro-poor”. However, the extent to which current deworming treatment programs achieve equitable coverage across wealth class and gender remains unclear, and the current public health metric of national deworming coverage does not include representation of inequity. This study develops a framework to measure both coverage and equity in global deworming to guide future programmatic evaluation, investment, and metric design.

**Methods:** We used nationally representative, geospatial household survey data that measured mother-reported deworming receipt in pre-school age children (age 1-4 years) in the previous 6 months. We estimated global deworming coverage disaggregated by geography, wealth quintile and gender and computed an equity index. We examined trends in coverage and equity index across countries, within countries, and over time. We used a regression model to compute the household correlates of deworming receipt and ecological correlates of equitable deworming.

**Findings:** Our study included 820,883 pre-school age children living in 50 STH-endemic countries between 2004 and 2017. Globally, the mean global deworming coverage in pre-school children was estimated at 36%. The sub-national coverage ranged from 0.5% to 87.5%, and within-country variation was greater than between-country variation in coverage. The equity index was undesirable (deworming was consistently concentrated in the wealthier populations) in every endemic region of 12 countries. Of the 31 study countries that WHO reported achieving the goal of 75% national coverage, 26 had persistent inequity in deworming as defined by the mean equity index. Deworming equity modestly improved over time, and within-country variation in inequity decreased over time. We did not detect differences in deworming equity by gender. We found the strongest household correlates of deworming to be vitamin A supplementation and receipt of three doses of diphtheria-tetanus-pertussis vaccine (DTP3), while the strongest ecological predictors of equitable deworming were regions with higher coverage of health services such as DTP3 and vitamin A supplementation.

**Interpretation:** Although mass deworming is considered to be “pro-poor”, we find substantial inequities by wealth, despite often high reported national coverage. These inequities appear to be geographically heterogeneous, modestly improving over time, and we found no evidence of gender differences in inequity. Future reporting of deworming coverage should consider disaggregation by geography, wealth, and gender with incorporation of an equity index to complement national deworming coverage.

**Funding:** Bill and Melinda Gates Foundation, Stanford University Medical Scientist Training Program

## Background

Soil-transmitted helminthiasis (STH) is the most prevalent neglected tropical disease of poverty with an estimated prevalence of 1.5 billion people.^1,2^ STH affects the most vulnerable people, helps to drives the cycle of poverty, and contributes to health inequities.^3,4^ STH is caused by infection with *Ascaris lumbricoides*, hookworm species of *Ancylostoma duodenale* and *Necator americanus*, and *Trichuris trichiura*, and is associated with a range of sequelae, including anemia, micronutrient deficiency, chronic abdominal pain, and stunting, some of which may be reversible with treatment.^5-8^ To address the public health burden of STH, the World Health Organization (WHO) recommends preventive chemotherapy (often referred to as mass “deworming”), which is a large-scale intervention that provides periodic administration of medications for empiric treatment of at-risk populations. In 2012, WHO broadened its school-based deworming program to include at-risk preschool-age children (1-4 years), in recognition of the significant disease burden in this age group that was otherwise not accessing treatment.^9^ During the past 15 years, deworming programs have made tremendous strides in scaling up coverage to reaching almost 600 million children annually, which makes it one of the largest public health programs in the world.^10,11^

While STH affects the most economically and socially disadvantaged, the degree to which current deworming treatment programs reach those most in need remains unclear. Equitable deworming would include all at-risk individuals, with coverage that is generally proportional to risk of STH and at least equal by economic and educational status, gender, and geographic zone. In some cases, equitable deworming may even be “pro-poor” given the higher risk of STH in more impoverished populations.^3,4^ Historically, the conventional metric for deworming has been national deworming coverage, which is used to track progress of country programs and to set the WHO goal of achieving 75% national coverage or greater by 2020.^4^ However, national deworming coverage does not necessarily reflect equity by economic status, gender, or geography.^12^ Specifically, national deworming coverage may be subject to “tyranny of averages” which can obscure low coverage in significant geographic regions, economically disadvantaged populations, or other key subgroups.^12^ As a result, our understanding of the key correlates of deworming receipt that may be related to inequity remains less clear.

While the goal of equity lies at the foundation of global health, emphasis on measuring and tracking equity in public health programs has increased with the Sustainable Development Goals.^13^ There is an abundant literature on simple metrics to estimate equity for various public health programs (e.g. concentration indices, ratios) and case examples on their use, such as with child mortality,^14,15^ yet their application to routine measurement and evaluation of programs in global health is limited. WHO recently provided guidance to public health programs on simple equity metrics and disaggregating programmatic data by geography, wealth, and gender.^15^

Mass deworming has been promoted as an example of an equitable and even “pro-poor” intervention in global health, yet the degree to which this is true remains unclear given limited programmatic monitoring of equity with disaggregation of data or incorporation of equity measures. To empirically test the degree to which deworming for STH is equitable and “pro-poor”, the aim of this study was to estimate the global status and trends in sub-national deworming coverage and equity by wealth and gender.

## Methods

We used nationally representative, geospatial household survey data from the Demographic and Health Surveys (DHS) to estimate global, national, and sub-national coverage disaggregated by wealth quintile and gender. We analyzed DHS data in pre-school age children (1-4 years) over the period 2004 to 2017. We computed equity indices for wealth by gender, and examined trends across space and time. We used a regression model to measure the household correlates of deworming receipt and ecologic correlates of deworming equity by region.

### Data and study measures

We used the DHS to examine global, national, and sub-national coverage and inequity in deworming, and the associated correlates of deworming. The DHS are nationally representative cross-sectional surveys on demographic, health, and health system indicators that are conducted in many low- and middle-income countries approximately every five years.^16^ We analyzed all available DHS surveys that included data on deworming in pre-school age children (1-4 years). We excluded data from countries non-endemic for STH or where deworming was not recommended due to low STH prevalence (see Appendix). For STH-endemic countries, we excluded geographic areas of each country that were non-endemic for STH or where deworming was not recommended due to low STH prevalence. This was determined using the WHO preventive chemotherapy databank, recent STH mapping data from ESPEN and other sources, and, in some cases, correspondence with national NTD program managers or WHO NTD focal points (see Appendix for details).^17,18^ Estimates of drug coverage and equity were based on the most recent DHS survey for each country. For analyses related to temporal trends or correlates of deworming, we included all surveys when multiple DHS surveys were available for a country. All DHS data are publicly available upon request.^16^ This study relied on secondary data that was not individually identifiable and did not require human subject research approval.

The key outcome variable was deworming receipt based on the mother’s report of whether her children received “drugs for intestinal parasites in the past 6 months” prior to the survey. We estimated coverage disaggregated by wealth quintile (a composite index of household assets, standardized to each survey), gender, and geography. We used DHS survey data on pre-specified variables to measure the correlates of deworming receipt, which included: child’s age and gender, mother’s age (<30 years or older), mother’s education (defined as no education, some primary school, and completion of primary school or higher education), wealth quintile, urban or rural residence (based on country definition), access to improved water and sanitation (binary designations based on definitions used by the UNICEF/WHO Joint Monitoring Program), and receipt of the third dose of diphtheria-tetanus-pertussis vaccine and vitamin A as markers of healthcare access.^19^ We excluded observations with missing data on relevant covariates.

### Statistical analysis of coverage and equity

We computed inequity in deworming receipt by wealth using an “equity index”, which was estimated using a concentration index. The concentration index is scalar measure of equality when considering a binary outcome (e.g. deworming receipt) over a range of another socioeconomic variable (e.g. composite wealth score).^20^ The generalized calculation involves the summation of binary deworming receipt weighted by wealth quintile (normalized to zero) over the population with adjustment for overall deworming coverage and population size.^20^ We used the Erreygers correction which provided numerous technical advantages over other concentration indices, including satisfying a mirror condition (inverse coding of the binary deworming variable would not affect the index).^20^ We computed the equity index at a sub-national level (DHS region) across all wealth quintiles. For interpretation, an equity index of 0 indicates that deworming was generally equally distributed across all wealth classes. A positive equity index (bounded to a maximum of 1) was “undesirable” and indicated that deworming was concentrated in children from wealthy families; the higher the value, the greater the concentration of deworming in children from wealthier families. A negative equity index (bounded to a minimum of −1) was “desirable” (preferentially “pro-poor” distribution) and indicated that deworming was concentrated in children from poorer families; the lower the value, the greater the concentration of deworming in children from poorer families. Importantly, the calculation of the equity index is related to both the degrees of equity and mean coverage of deworming, where decreasing equity (deworming concentrated in wealthier populations) and increasing coverage are positively correlated with the index, although the calculation is more strongly driven by equity than mean coverage. We estimated the proportion of variance in concentration index explained by within country compared to between country variation, to understand whether inequity was driven more by national policy or within country differences. To estimate the proportion of variation explained between country differences, we performed a regression of the deworming receipt variable with country indicator variables and estimated the R^2^ value, whereas the remainder were attributable to within country differences.

### Statistical analysis of correlates of deworming receipt and equity

We computed the household-level correlates of deworming using all available survey data, including multiple surveys for a given country. The analysis used a multivariable logistic regression model using a dependent variable of mother-reported deworming receipt to compute the correlates of deworming.^21^ The analysis included a pre-specified set of correlates based on prior literature as described above.^6^ We included country fixed effects, which controlled for all time-invariant differences between countries (e.g. baseline country differences in economic status or STH disease burden). We estimated the marginal effects for correlates of deworming receipt among children for ease of interpretation, and used robust standard errors clustered by country and survey.^21^

We computed the ecological correlates of equity in deworming using all available survey data averaged at a regional level. We used a ordinary least squares regression model using a dependent variable of region-level equity index, and independent variables of region-level averages of the pre-specified set of correlates as described above. We estimated the absolute change in concentration index associated with a unit value for each covariate at an ecological region level.

## Results

### Study descriptive analysis

The study included data on 820,883 pre-school age children over the period of 2004 to 2017, with the most recent surveys (defined as only the most recent survey for each country) with data from 628,489 children. We included 77 DHS surveys from 50 countries, including from Africa (n = 35), the Americas (n = 5), Asia (n = 9), and Europe (n = 1) (Table 1). Globally, our study population of pre-school children was 49% female; 72% lived in rural settings, 47% had access to improved toilets, 64% had access to improved drinking water, 29% had mothers with greater than primary level education, and 66% received their third dose of diphtheria-tetanus-pertussis immunization (see Table S1 in Appendix).

**Table 1:**
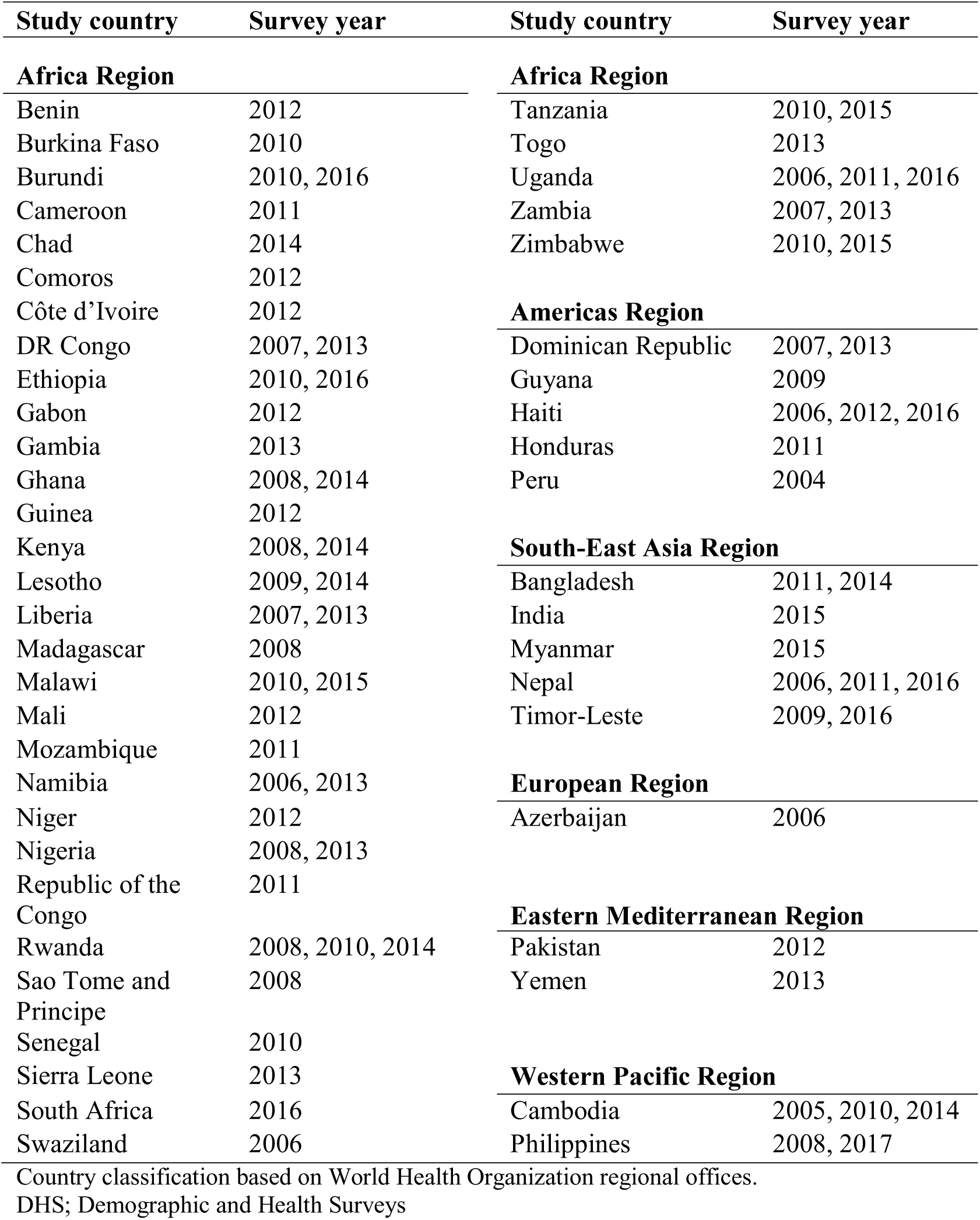
Study countries and DHS surveys.

### Global, national, and sub-national coverage and inequity in deworming

Across all countries and years, 36% of mothers reported that their child received deworming in the past 6 months. Deworming coverage varied substantially across countries, with an even higher degree of within-country variation (Figure 1). The coverage of deworming in endemic areas varied by country, from 5.1% (Azerbaijan) to 73.6% (Rwanda) and varied further at the sub-national or regional level, which ranged from 0.5% (Zamfara, Nigeria) to 87.5% (Kavango, Namibia). The estimated proportion of variation in deworming coverage explained by within-country factors was 91%, while the remaining 9% was attributable to between-country factors (e.g. national policies and programs).

**Figure 1:**
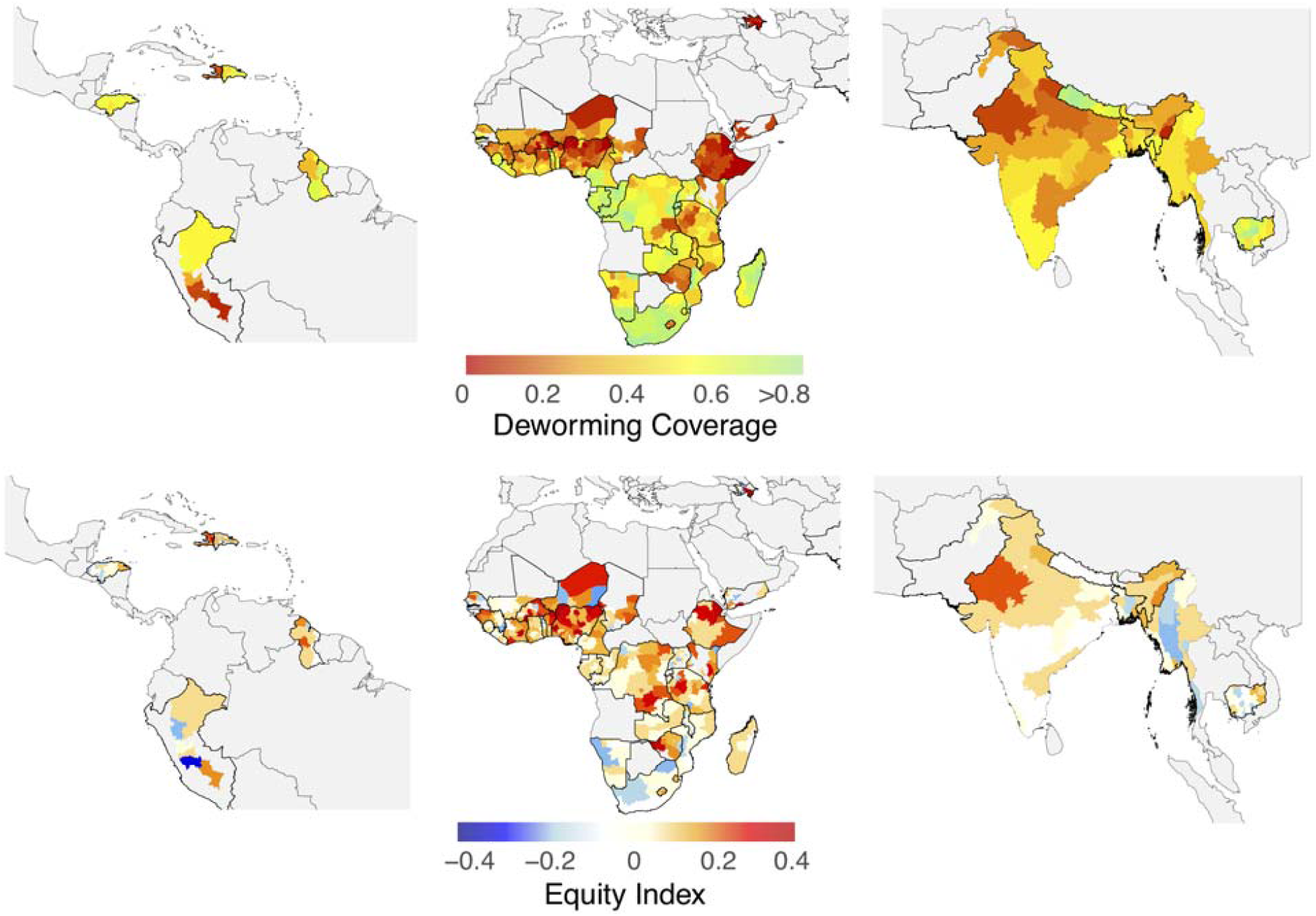
Global deworming coverage and inequity. We estimated sub-national deworming coverage (top panel) and inequity measured in the equity index (bottom panel) for all study countries. We used geospatial household survey data from the Demographic and Health Survey from 2004 to 2017. The map was geographically divided into the Americas (left column), Africa (middle column), and Asia (right column) for data visualization purposes. For the interpretation of the equity index, a positive index was undesirable and indicated deworming was concentrated in wealthier households, while a negative index was desirable and indicated deworming was concentrated in poorer households.

At the global level, deworming of pre-school children in the previous six months increased linearly with household wealth, ranging from deworming coverage of 27.5% for children in the lowest wealth quintile to 41.5% in the highest quintile. Of the 50 countries, 12 had an undesirable equity index in every endemic region of the country. Of these, Nigeria, Haiti, Burkina Faso, Azerbaijan, and Ethiopia had the highest level of deworming inequity; for Nigeria this inequity translated to a 28% absolute difference in deworming coverage between children in the lowest and highest wealth quintile. A few countries including Cambodia, Malawi, Honduras, Namibia, South Africa, Rwanda, Nepal, Peru, and Gambia had relatively equitable deworming across regions of their country (i.e. mean equity index that approximated zero). Notably, some countries had substantial within-country variation in the equity index, including Azerbaijan, Burkina Faso, India, Nigeria, and Philippines. Of the 31 study countries that WHO report to have achieved a national deworming coverage greater than the public health goal of 75%, we found 26 had persistent inequity by wealth in majority of the country (defined as mean equity index greater than zero). We calculated equity index independently by gender, and notably did not find substantial systematic differences between boys and girls in deworming coverage equity across study countries (Figure 2).

**Figure 2:**
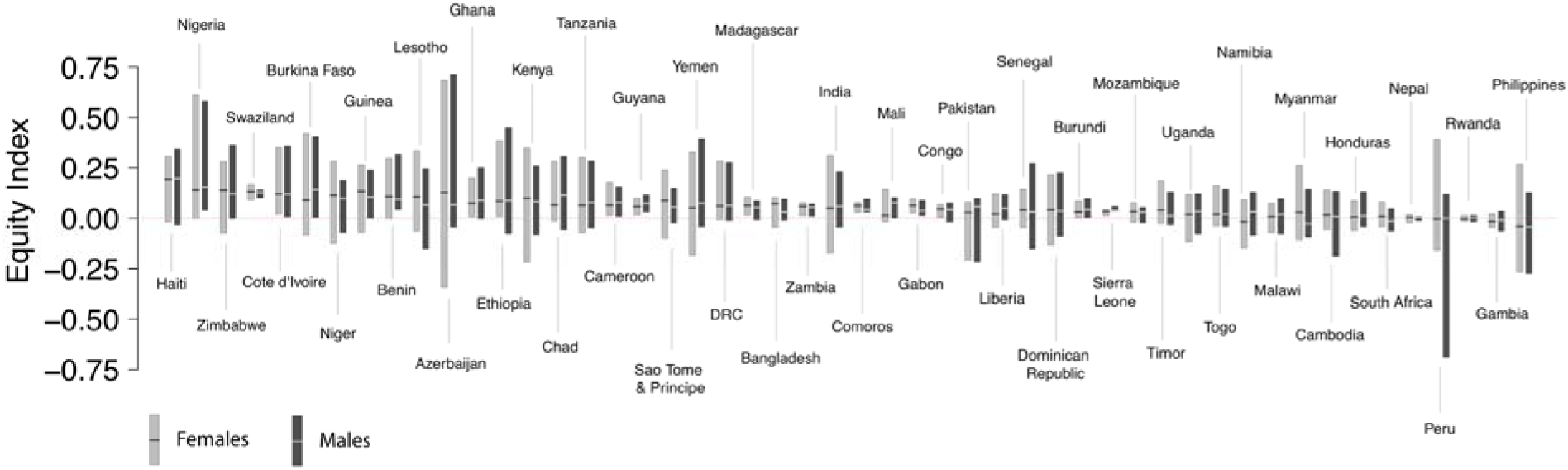
Geographic variation in deworming equity by country and gender. We estimated inequity (measured in the equity index) in deworming at a sub-national level for each study country, which provided data on within country variation of deworming inequity. The height of each box provides the 2.5^th^ and 97.5^th^ percentile of the equity index computed for each country. The analysis was conducted independently by gender (light bar-female, dark bar-male) for comparison of inequity by gender for each study country. For interpretation, the height of each bar corresponds with the magnitude of within country variation in inequity; where a positive index was undesirable and indicated deworming was concentrated in wealthier households, while a negative index was desirable and indicated deworming was concentrated in poorer households.

For countries with multiple DHS surveys, we found modest improvements in deworming coverage and equity over time, meaning more desirable mean equity index or reduced within-country variation in deworming equity. We found an average increase in national deworming coverage by 0.7% annually. For temporal trends in equity, we estimated that seven countries had notable improvements to deworming equity, 13 countries had no changes, and 2 countries became less equitable (see Table S2 in Appendix). We correlated the regional mean deworming coverage and equity index, and found that regions with higher coverage had more equitable deworming distribution (Figure 3). When estimated sub-national deworming coverage approached 70% using study coverage estimates, the average equity index at the sub-national level approached zero, suggesting equitable distribution of deworming. Similarly, when we estimated how sub-national changes in mean deworming coverage were associated with corresponding sub-national changes in equity index for a given country, we found that higher coverage was related to more desirable equity index, although inequities still persisted (Figure 3).

**Figure 3:**
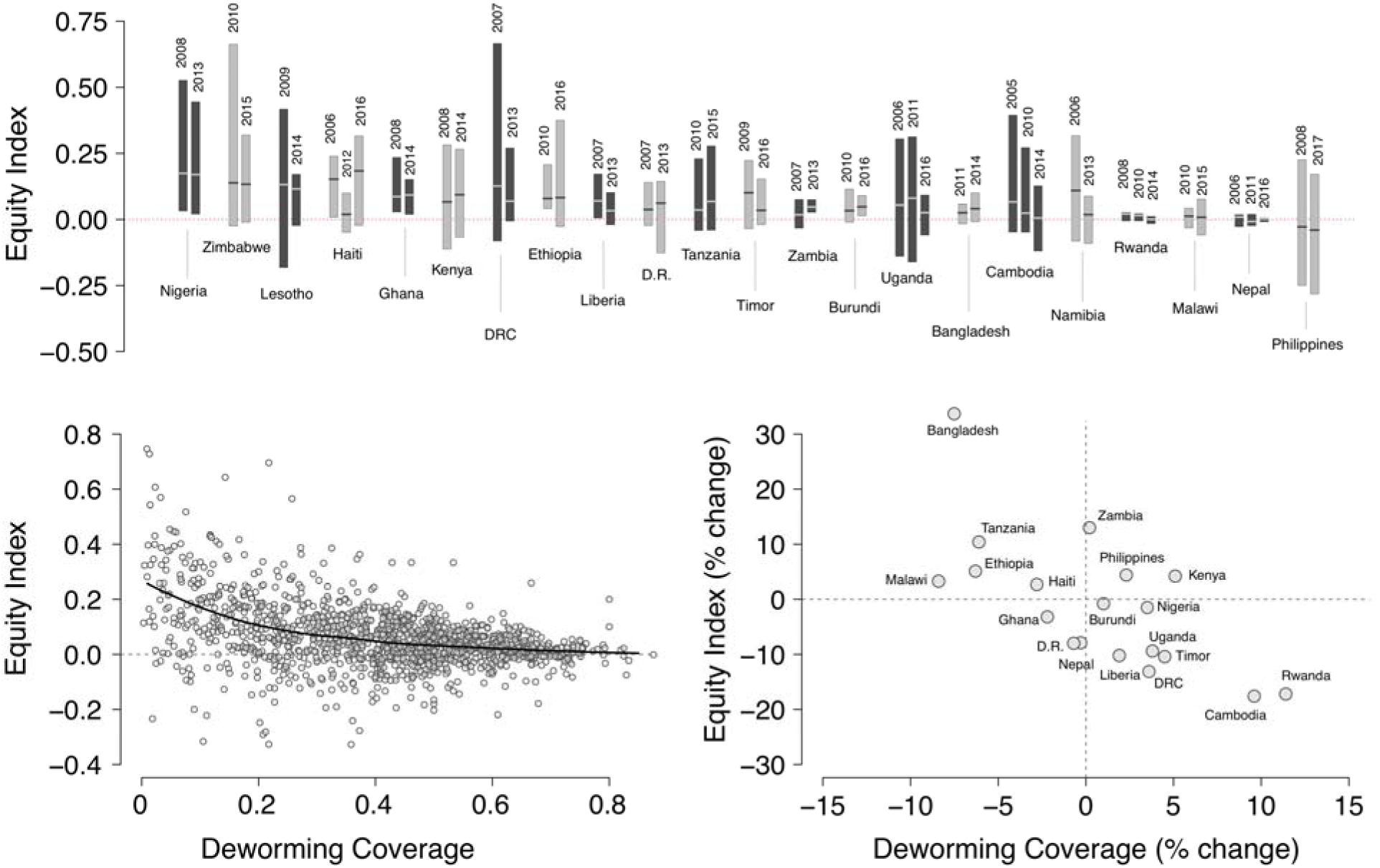
Trends in deworming inequity over time and by programmatic coverage. (Top panel) We estimated inequity (measured in the equity index) for deworming in countries with surveys at different time periods, which provided data on country trends for geographic variation in deworming inequity. The height of each box provides the 2.5^th^ and 97.5^th^ percentile of the equity index computed for each country. The bars were grouped by country, with more recent surveys shown rightward. For interpretation, deworming inequity can improve for a country with reduced height of the bar (reduced within country variation in inequity) or lower mean for equity index (overall improvement in equity). (Bottom left panel) We correlated the sub-national mean deworming coverage and equity index for all study countries. (Bottom right panel) We estimated the annualized change in mean deworming coverage and the corresponding annualized change in equity index for study countries with surveys at different time periods.

### Correlates of deworming receipt and equity

In a multivariate analysis, we found that receipt of deworming medicine during the previous 6 months was most strongly associated with receipt of vitamin A supplementation and three doses of diphtheria-tetanus-pertussis vaccine, and maternal education (Table 2). We found weaker associations between deworming receipt and older child age, greater family wealth, and improved drinking water. There was a modest, borderline relation between deworming receipt and gender (Adjusted marginal effects: −0.005, 95% CI: −0.009, −0.001), which corresponded with a 0.5% coverage difference by gender. In an ecological multivariate analysis, we found equitable distribution of deworming was most strongly associated with high coverage of three doses of diphtheria-tetanus-pertussis vaccine and vitamin A supplementation, average child age, and wealthier populations at a region level (Table S4 in Appendix). The estimated explanatory value of gender on the variation in deworming equity was <1%.

**Table 2:** Regression model estimating the correlates of deworming receipt in pre-school age children.

| **Variable** | **Adjusted**<br>**marginal effects** | **95% CI** |
| --- | --- | --- |
| Age (year) | **8·7** | **(7, 10·5)** |
| Gender (female) | **-0·5** | **(-0·9, -0·1)** |
| Wealth quintile | **1·5** | **(0·5, 2·5)** |
| Mother's age (<30 years) | **-1·2** | **(-1·8, -0·5)** |
| Rural residence | 0·7 | (-0·6, 2·1) |
| Mother's education (tertile) | **4** | **(2·6, 5·4)** |
| Improved drinking water | 1·2 | (0, 2·5) |
| Improved toilet facility | 1·2 | (-0·1, 2·5) |
| Receipt of 3<sup>rd</sup> dose of DPT vaccine | **13·3** | **(11·6, 15·1)** |
| Receipt of vitamin A | **34·3** | **(30·1, 38·5)** |

Bolded findings indicate a relationship with a $p < 0.05$ . Marginal effects refer to absolute change in mean deworming coverage associated with one unit change in variable. Robust standard errors clustered by country and survey.

When considering healthcare access, we found that of the 66% of pre-school children who completed their third dose of diphtheria-tetanus-pertussis vaccine, only 42% of these children had received deworming in the previous 6 months; deworming coverage was estimated at 21% for children who did not receive this vaccine. Similarly, of the 49% of pre-school children who had received vitamin A supplementation in the previous 6 months, 48% of these children had also been dewormed.

## Discussion

In this study, we applied a geospatial empirical analysis to household survey data from over 820,000 pre-school age children in 50 countries to estimate the global status and trends in coverage and equity for deworming programs. We find that while NTDs and deworming are considered “pro-poor” interventions, in most countries, deworming coverage is highly geographically variable and generally not equitable despite often reporting a high national coverage. In the 31 study countries that reported to WHO a greater than 75% national coverage, 26 had persistent inequity in this analysis. Notably, we do not find systematic evidence of gender-related differences in inequity. Over time, inequities in deworming by wealth seem to be modestly improving with associated reductions in within country variation in inequity. We found the strongest correlate of deworming receipt as well as equitable deworming in a region to be receipt of vitamin A supplementation and receipt of three doses of diphtheria-tetanus-pertussis vaccine. The findings from our global empirical analysis support the need for routine measurement of equity in national deworming programs, and potential to incorporate secondary data sources into the measurement framework.

The goal of equity has been at the foundation of global health, despite the challenge of providing public health programs equitably across geography, wealth classes, gender, and race.^22^ More recently, equity has been more formally considered in the measurement and evaluation of global health programs. Goal 10 of the Sustainable Development Goals is to reduce inequities, and WHO now includes equity as a core consideration in their evaluation of evidence and guidelines.^13^ Many recent analyses have addressed equity in global health programs, such as in the case of diphtheria-tetanus-pertussis vaccination where data illustrated a persistent but improving degree of inequity.^23^ Notably, recent work created a multi-disease NTD index that represented a composite estimate of coverage of multiple NTD programs across a country.^24^ However, while these analyses have explored equity, most have only examined coverage trends at a national level rather than equity at a sub-national level and lack disaggregation by wealth and gender, which limits interpretation and development of a simple equity metric to incorporate into programmatic use.

Historically, the key metric in deworming for defining public health goals and monitoring programmatic progress has been national coverage. By 2020, WHO defines the public health goal for STH as achieving greater than 75% national coverage of deworming in pre-school children. However, overall national coverage estimates can appear high while neglecting significant inequities across geography, wealth, and gender, which has been referred to as the “tyranny of averages.”^12^ Indeed, of the 31 study countries that WHO reported achieving the public health goal of 75% coverage, 26 had persistent inequity based on the equity index (as well as low coverage in regions of the country) as well as lower deworming coverage when estimated with study data. Many countries such as Burkina Faso and Zimbabwe have substantial variation in within-country deworming equity, which is not captured with national coverage alone. We estimated that 91% of sub-national variation in deworming was explained by within-country (rather than between-country) factors; the largest variations in coverage are not explained at a national level (e.g. country guidelines, national policy), underscoring the importance of reporting sub-national coverage data.

This analysis incorporated a metric of an equity index and computed correlates of deworming to complement reporting of programmatic coverage alone. We found that substantial inequity in deworming persists, with large within-country variation, which appears to be modestly improved over time. Notably, there were not clear differences in equity by gender. However, this does not exclude gender inequity across many settings and supports the need for ongoing measurement of gender equity. We also estimated that the strongest global correlate of deworming to be receipt of vitamin A supplementation and receipt of three doses of diphtheria-tetanus-pertussis vaccine, and maternal education. These relationships likely related to the common co-administration of deworming and vitamin A in this age group, access to general health services (both household and systems-level indicator), and access to health information given a mother’s education level. The strongest ecological predictors of equitable deworming shared these characteristics, and were identified as regions with higher coverage of DTP3 and vitamin A supplementation. The data suggest that where strong healthcare services exist, such as routine healthcare (e.g. vaccination) and public health programs (vitamin A provision), deworming equity is likely to be greater. The use of DHS survey data also identified the potential to integrate deworming efforts with other interventions (e.g. vitamin A) that have achieved high coverage where deworming coverage is persistently inequitable or low.

This study proposes a new analytical framework to formally incorporate equity into routine evaluation of deworming programs that can complement the current metric of national deworming coverage. Through incorporation of existing secondary datasets from the DHS, this approach provides disaggregated sub-national data and an equity index that can potentially mitigate ongoing inequities if used to guide programmatic decisions that respond to inequities. To fully realize the potential of this framework, the collaborative support of WHO, Ministries of Health, and non-governmental organizations are necessary as well as consideration of other public health interventions to evaluate the comprehensive needs of countries.

The findings of this study should be interpreted within the context of the limitations of data and study design. The main outcome variable of deworming receipt in past six months was proxy-reported by the mother and subject to recall bias, which could in theory be differential by wealth status, although probably less likely by gender. Notably, data suggest that the reliability of self-reported and mother proxy-report for deworming medicines is reasonably good for mass drug administration.^25^ The proxy-reported deworming receipt variable in this study would have included both programmatic deworming (e.g. preventive chemotherapy, through child health days) and unprogrammed deworming (e.g. local pharmacy, healthcare center).^26,27^ However, since deworming is commonly given with vitamin A supplements during child health days; the fact that 82% of children dewormed during the previous 6 months also were reported to have received vitamin A supplements during the same timeframe suggests that the majority of deworming occurred in the context of preventive chemotherapy. Importantly, wealthier households may be more likely to purchase or obtain deworming medication in an unprogrammed setting (e.g. health clinic, pharmacy), which would bias study findings towards an inequitable distribution. The choice of alternative formulations for the equity index affects the mean estimate, and thus the relative comparisons for trends are more robust than absolute value. While the study measured equity in deworming, we believe this study has implications for many preventive chemotherapy programs for NTDs (e.g. lymphatic filariasis, schistosomiasis), although we are unable to generalize to NTDs that use different drug delivery platforms (e.g., community-based or school-based drug distribution). We excluded regions known to be non-endemic for STH or sufficiently low prevalence where preventive chemotherapy is not recommended, although up-to-date data on STH prevalence and geographical coverage of national deworming programs were not always complete (see Appendix).

This study found that provision of deworming in pre-school age children is highly variable and generally not reaching the poor equitably. In the context of continued scale up of deworming programs in which the primary metric is national coverage, we provide a new analytical framework using household survey data, geospatial analysis, and equity index to inform future tracking of the equity of programmatic deworming to address the burden of STH and other NTDs.

## Authorship contribution

The corresponding author had full access to all of the data in the study and takes responsibility for the integrity of the data and the accuracy of the data analysis.

Study design- NCL, EB, and DA

Data analysis- NCL, SHN

Data interpretation- All authors

Wrote first draft- NCL

Contributed intellectual content and approved final draft-All authors

## Role of the Funding Organization or Sponsor

The funding organizations had no role in the design and conduct of the study; collection, management, analysis, and interpretation of the data; and preparation, review, or approval of the manuscript; or the decision to submit the manuscript for publication.

## Previous presentations

This work was presented by NCL as an oral abstract at the American Society of Tropical Medicine and Hygiene (New Orleans, USA 2018) and at the WHO 10^th^ annual measurement and evaluation working group meeting on neglected tropical diseases (Geneva, Switzerland 2019).

